# Microglia remodel synapses by presynaptic trogocytosis and spine head filopodia induction

**DOI:** 10.1101/190975

**Authors:** Laetitia Weinhard, Urte Neniskyte, Giulia di Bartolomei, Giulia Bolasco, Pedro Machado, Nicole L. Schieber, Melanie Exiga, Auguste Vadisiute, Angelo Raggioli, Andreas Schertel, Yannick Schwab, Cornelius T. Gross

## Abstract

Microglia are highly motile glial cells that are proposed to mediate synaptic pruning during neuronal circuit formation. Disruption of signaling between microglia and neurons leads to an excess of immature synaptic connections, thought to be the result of impaired phagocytosis of synapses by microglia. However, until now the direct phagocytosis of synapses by microglia has not been reported and fundamental questions remain about the precise synaptic structures and phagocytic mechanisms involved. Here we used light sheet fluorescence microscopy to follow microglia-synapse interactions in developing organotypic hippocampal cultures, complemented by three-dimensional ultrastructural characterization using correlative light and electron microscopy (CLEM). Our findings define a set of dynamic microglia-synapse interactions, including the selective partial phagocytosis, or trogocytosis (*trogo-*: nibble), of presynaptic structures and the induction of postsynaptic spine head filopodia by microglia. These findings allow us to propose a mechanism for the facilitatory role of microglia in synaptic circuits remodeling and maturation.

## Introduction

Microglia are glial cells that derive from the myeloid hematopoietic lineage and take up long-lived residence in the developing brain. In response to brain injury microglia migrate to the site of damage and participate in the phagocytic removal of cellular debris^1^. The recent discovery that microglia are also highly motile in the uninjured brain^2,3^, continuously extending and retracting processes through the extracellular space, suggests that they may participate in brain homeostasis. Clues to this activity come from observations that during early postnatal development microglia undergo morphological maturation that matches synaptic maturation^4^ and that they express receptors for neuronal signaling factors that are upregulated during this period^5^. These data, combined with the known phagocytic capacity of myeloid cells, led to the hypothesis that microglia may have a role in the phagocytic elimination of synapses as part of the widespread pruning of exuberant synaptic connections during development^6,7^. This hypothesis was supported by two studies that reported the selective engulfment of synaptic structures by microglia and the appearance of excess immature synapses in mice lacking either the fractalkine^8^ (Cx3cl1/Cx3cr1) or complement component^9^ (C1q/C3/CR3) microglia signaling pathways.

Numerous studies have confirmed an important role for microglia in promoting synapse and circuit maturation. Knockout mice lacking complement factors show cortical excitatory hyperconnectivity^10^ supporting a role for microglia in the elimination of excess synapses in the mammalian neocortex. Knockout mice lacking fractalkine receptor show transient excitatory hyperconnectivity followed by weak synaptic multiplicity and reduced functional brain connectivity in adulthood^11,12^ suggesting that a failure to eliminate synapses at the correct developmental time prevents the normal strengthening of synaptic connections. Notably, inhibitory synapses appear to be unaffected by disruptions of neuron-microglia signaling^11^. Microglia are also likely to be required for environment-induced brain plasticity as mice lacking microglia P2Y_12_ receptors show deficits in early monocular deprivation-associated visual cortical plasticity^13^. At the same time, studies have pointed to a role for microglia in synapse formation in the adult brain, showing that they can elicit calcium transients and the formation of filopodia from dendritic branches^14^ and that they are required for learning-induced synapse formation^15^. Together, these studies suggest that microglia have a complex role in shaping maturing circuits.

An important question raised by these studies is whether synaptic phagocytosis by microglia underlies some or all of these phenotypes. Unfortunately, support for a role of microglia in the phagocytic elimination of synapses is based entirely on indirect evidence - localization of synaptic material within microglia in fixed specimens and increased synapse density following the disruption of microglial function.

In this study we set out to test the hypothesis that microglia engulf and eliminate synapses during mouse hippocampal development. First, we carried out quantitative confocal microscopy analysis of microglia-synapse interactions in fixed hippocampal tissue. Microglia were found to contact dendritic spines, but contrary to previous reports, we found no evidence for the elimination of postsynaptic material. Second, we confirmed these data at the ultrastructural level by correlative light and electron microscopy (CLEM) using focused ion beam scanning electron microscopy (FIB-SEM) and discovered evidence for the partial phagocytosis, or trogocytosis, of presynaptic material by microglia. Third, we carried out time-lapse light sheet microscopy of microglia-synapse interactions in organotypic hippocampal cultures and confirmed the exclusive trogocytosis of presynaptic material. Time-lapse microscopy also revealed the frequent induction of spine head filopodia at sites of microglia-synapse contacts and these were confirmed at the ultrastructural level. Our findings provide the first direct evidence for the phagocytic elimination of synapses by microglia in living brain tissue, and suggest that microglia facilitate synaptic circuit maturation by a combination of trogocytosis of presynaptic material and the remodeling of postsynaptic sites.

## Results

### No evidence for phagocytic elimination of spines by microglia

To identify the period of maximal microglia phagocytic activity in postnatal hippocampal development we performed immunocolocalization analysis of Iba1-labelled microglia with CD68, a phagosomal marker, in fixed sections at postnatal day 8 (P8), P15, P28, and P40 (Supplementary Fig. 1a,b). CD68 immunoreactivity was high at P8-P15 with a peak at P15, and a gradual decrease in the following weeks. The CD68 immunoreactivity pattern at P15 was confirmed in genetically labelled microglia (Supplementary Fig. 1c,d). These findings indicate that the second postnatal week of mouse hippocampal development is likely to be a period of active microglia phagocytosis, and suggested that this period may be most relevant to search for evidence of phagocytic elimination of synapses by microglia.

Previous studies have presented indirect evidence for the phagocytic engulfment of dendritic spines in hippocampus and visual cortex^8,13,16^. For example, immunoreactivity for postsynaptic density 95 (PSD95) protein was found to be localized inside microglia by confocal, super-resolution, and electron microscopy^8^. We attempted to confirm these findings using cytoplasmic neuronal and microglia markers, in triple transgenic mice expressing GFP in sparse excitatory neurons (*Thy1*::EGFP^17^), and tdTomato in microglia (*Cx3cr1*::CreER^15^; *RC*::LSL-tdTomato^18^). We analyzed over 8900 spines from secondary dendrites of GFP+ neurons in the CA1 *stratum radiatum* of fixed hippocampal tissue at P15 and found that about 3% of spines were contacted by microglia (n = 294). The majority of these contacts presented a relatively minor colocalization of spine and microglia fluorescence (apposition, Fig. 1a,c; n = 171, 1.9% of spines), while others were characterized by more extensive colocalization as defined by more than 70% of the spine surface contacted by microglia (encapsulation, Fig. 1b,c; n = 123, 1.4% of spines). For each encapsulation event we carefully examined the integrity of the spine head, neck, and dendritic shaft and found that in all cases, even in those where the entire spine surface appeared to be contacted by microglia, an intact GFP+ spine neck remained visible (Figure 1b-d). Thus, using cytoplasmic markers to visualize neuronal and microglia material we were not able to confirm earlier evidence suggesting the phagocytic engulfment of dendritic spines. To explore whether phagocytic activity might nevertheless be associated with microglia-spine contacts, we performed immunofluorescence colocalization of the phagosomal marker CD68 and microglia-spine contacts. Approximately 15% of microglia-spine contacts (apposition or encapsulation) showed apposed CD68 immunoreactivity (Fig. 1d) suggesting an involvement of local phagocytic activity in the microglia-neuron contact event.

**Figure 1.**
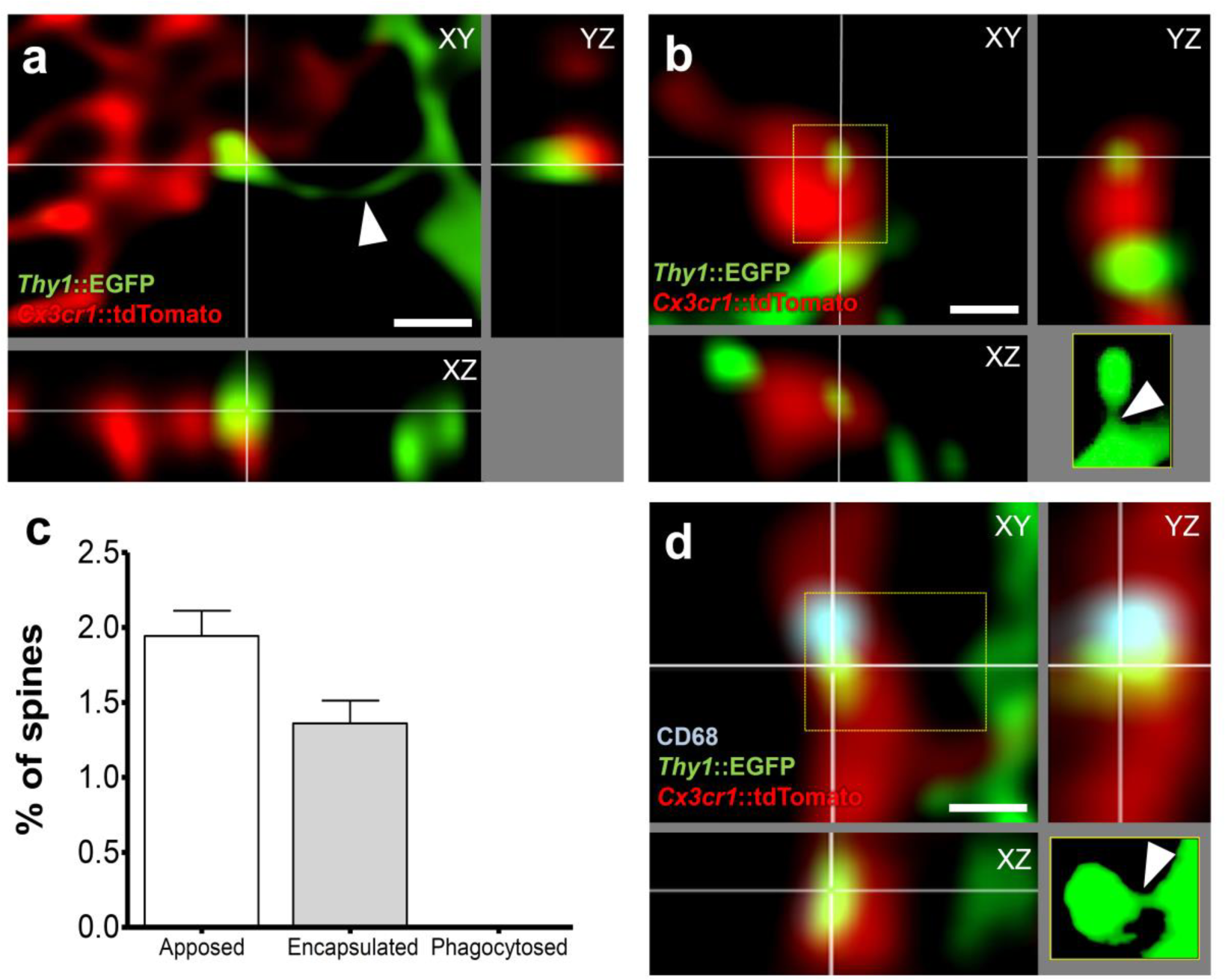
Microglia do not phagocytose dendritic spines. Representative images of microglia (red, *Cx3cr1v*.CreER; *RC*::LSL-tdTomato) **(a)** apposing or **(b)** encapsulating a dendritic spine (green, *Thy1::*EGFP; yellow insert: projection of the undeconvolved z-stack containing the contacted spine). Note that the neck of the contacted spine is intact (white arrow head), **(c)** Quantification of microglia-spine contacts. No spine was found phagocytosed, as contacted spines were always attached to their dendrite, **(d)** Encapsulated spine localizing next to a phagocytic compartment (blue, CD68 immunostaining, scale bar = 0.5 pm).

To test this hypothesis we developed a correlative light-electron microscopy (CLEM) approach based on previously published methods^20,21^ to first identify rare microglia-spine contact events by fluorescence microscopy and then reconstruct in three-dimensions the surrounding ultrastructure by electron microscopy (Fig. 2a, Supplementary movie 1). Following fixation of hippocampal tissue in a manner compatible with electron microscopy, interaction events were identified by confocal microscopy. Once a region of interest (ROI) was identified, concentric brands were etched into the surface of the fixed tissue with a UV-laser microdissector microscope and the tissue was embedded and prepared for electron microscopy. The resulting trimmed block of tissue was then subjected to X-ray imaging to identify vasculature and nuclear landmarks and align these with the branding. The block was then subjected to focused ion beam scanning electron microscopy (FIB-SEM) at 5 nm lateral pixel size and 8 nm section thickness using the landmarks as guides to capture the ROI. Four ultrastructural image stacks (up to 35 x 20 μm area; 2300 images) were obtained containing a total of 8 apposition and 5 encapsulation ROIs. Three-dimensional ultrastructural reconstruction confirmed microglia-spine apposition events (7/8 apposed; Fig. 2f). However, most encapsulations appeared as simple appositions (3/5), with only one ROI showing more than 50% of the spine contacted by microglia (Figure 2b-f). These findings show that caution must be exercised when interpreting colocalization of synaptic material with microglia using light microscopy. Moreover, these data do not support the hypothesis that microglia phagocytose dendritic spine material.

**Figure 2.**
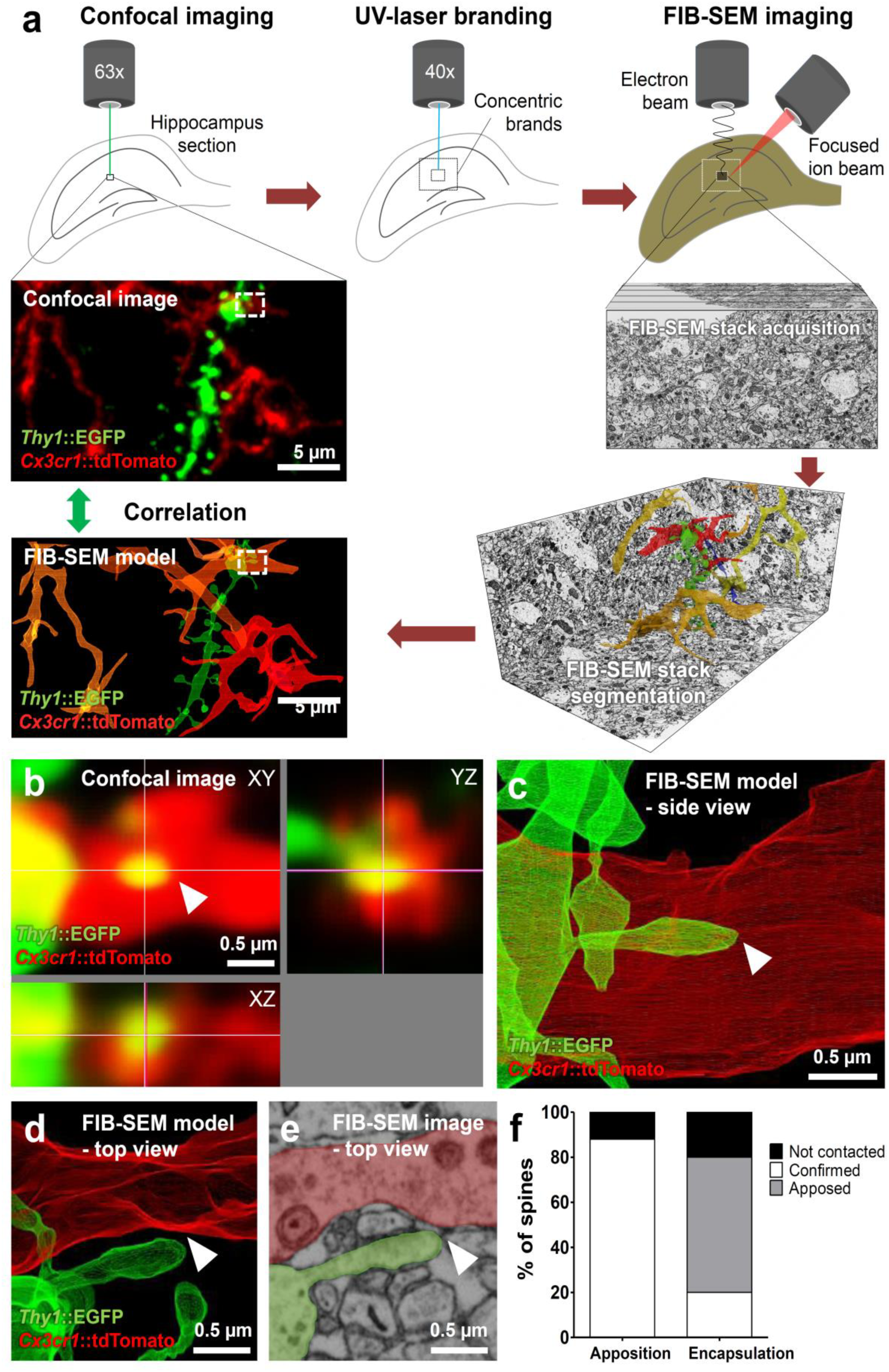
CLEM analysis of microglia-spine interactions. **(a)** Schematic of Correlative Light and Electron Microscopy (CLEM) workflow, **(b)** Confocal orthogonal view of a region of interest (ROI, dotted line in [a]) containing a spine encapsulated by microglia, **(c)** Segmentation of the ROI containing the encapsulation from the corresponding electron microscopy dataset, side view, **(d)** Top view of the ROI revealed that the spine was not encapsulated, with **(e)** no sign of elimination, **(f)** Quantification showed that the majority of the encapsulations observed by confocal microscopy were simple appositions by electron microscopy.

### Microglia trogocytose presynaptic elements

Previous studies have argued for the phagocytosis of presynaptic material by microglia^8,9,13,16^ and we re-examined our electron microscopy datasets for evidence of microglia engulfment of identified presynaptic structures. We examined over 50 μm^3^ of microglial material for a total surface of 560 μm^2^ from 8 reconstructed cells. We found 17 confirmed double-membrane inclusion bodies (Fig. 3d). Two of these contained putative presynaptic vesicles (Fig. 3a,d, Supplementary movie 2) suggesting that presynaptic material is a substrate for microglia phagocytosis. In addition, we found 20 double membrane structures that appeared to be in the process of engulfment. Many involved axonal shafts (8/20; Fig. 3b,d, Supplementary movie 3) and a smaller number involved synaptic boutons (3/20). We also frequently observed microglia autophagocytosis in which microglial processes were being engulfed by the same cell (9/20; Fig. 3c,d). Analysis of the size distribution of inclusions revealed that the presynaptic material engulfed from boutons and axons typically ranged between 0.01 and 0.05 μm^3^ (Fig. 3e), with an average diameter of 253 ± 24 nm. This shows that presynaptic structures are not entirely phagocytosed by microglia but rather “trogocytosed”, a term originally coined to describe membrane transfer in immune cells^22^ and later extended to refer to partial phagocytosis by various cell types, including macrophages^23-25^. We also observed numerous invaginations of microglia facing boutons or axons that allowed us to reconstruct the putative sequence of events leading to the microglial digestion of these structures (Fig. 3f). Engulfment did not appear to be mediated by the formation of phagocytic cups as no microglial pseudopodia were observed at the contact site. Instead, boutons or axonal pinches appear to sink into microglial cytoplasm before closure of the membrane and subsequent trafficking. These findings argue for the specific trogocytosis of presynaptic structures by microglia and suggest that this activity may be oriented indiscriminately toward axons and synaptic boutons rather than selectively targeting the presynaptic active zone. To explore the dynamics of interactions between synapses and microglia we developed a time-lapse fluorescence imaging method in brain explant cultures (Supplementary Fig. 2a). Organotypic hippocampal slice cultures are known to undergo key developmental steps similar to those observed *in vivo*, including synapse maturation^26–29^, and have been previously used to study ramified-microglia function^30^. As shown previously, microglia initially respond to culturing by retracting their processes and assuming an activated phenotype^31^ (Supplementary Fig. 2b). Following one week in culture, however, microglia morphology resembles that found *in vivo*^32^ (Supplementary Fig. 2b,c). Time-lapse imaging of hippocampal cultures was performed using light sheet fluorescence microscopy in order to minimize light toxicity common to point-source beam scanning microscopes and to allow for the visualization of multiple fluorophores across very large fields of view (up to 0.5 x 0.5 x 0.2 mm) at relatively high frame rates (up to 1 frame/45 seconds) for protracted periods (up to 3 hours). To validate the technique and determine whether microglia in hippocampal explants showed *in* vivo-like physiology, we quantified the number and speed of process extension and retraction events (Supplementary Fig. 2d,e, Supplementary movie 4). No significant difference was observed between extension and retraction over time, with an average of 26 extension and 21 retraction events per cell per minute (Supplementary Fig. 2d, n = 4 cells, two-way ANOVA, main effect of time: F_2_, _12_ = 0.22, p = 0.81; main effect of direction: F_1_, _12_ = 0.96, p = 0.37). The speed of extension and retraction events was similar and stable over time (1.9 μm/min and 1.8 μm/min, respectively; Supplementary Fig. 2e) and consistent with previous *in vivo* imaging studies^3^.

**Figure 3.**
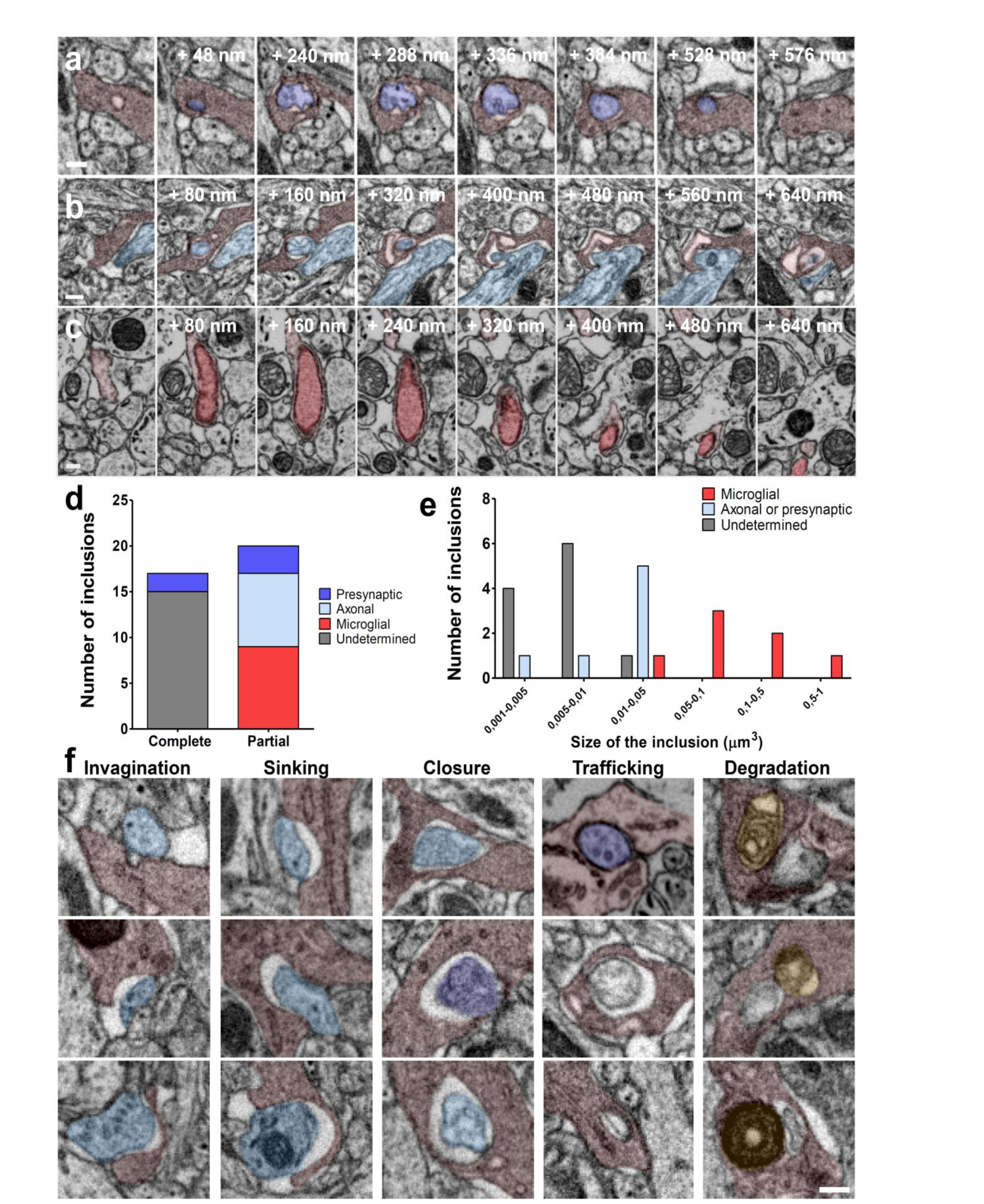
Microglia trogocytosis of presynaptic material. Representative FIB-SEM image sequences of **(a)** complete presynaptic inclusion containing 40 nm presynaptic vesicles (dark purple) inside microglia (red), **(b)** partial presynaptic inclusion containing axonal material (clear blue) inside a microglia, and **(c)** partial inclusion of a microglial process (dark red) inside microglia (clear red), **(d)** Quantification of microglial partial and complete inclusions, **(e)** Distribution of the volume of microglial inclusions, **(f)** Putative sequence of events leading to presynaptic material digestion by microglia, represented by a collection of 3 examples for each step (yellow: lysosomes, scale bar = 200 nm).

Next, we labeled presynaptic CA3 to CA1 Schaffer collateral projections with cytoplasmic near infra-red fluorescent protein (iRFP) following local adeno-associated viral infection (AAV-*Syn*::iRFP, Fig. 4a) of the CA3 region of organotypic slices from *Thy1*::EGFP; *Cx3cr1*::CreER; *RC*::LSL-tdTomato triple transgenic mice shortly after culturing. Importantly, microglia at the imaging site did not show any detectable morphological changes following viral infection as they were imaged two weeks later and 500 μm distant from the infection site. Consistent with our fixed electron microscopy data, we found clear evidence for the phagocytic engulfment of presynaptic material (Fig. 4b,d, Supplementary movie 5). Surprisingly, presynaptic engulfment events (n = 11 from 8 microglia analyzed) were rapid, frequently occurring in less than 3 minutes (Fig. 4c), which also raised the possibility that some phagocytic events occurring within our frame rate (1 frame/90 sec) went unnoticed. Notably, even in those cases where microglia were seen to phagocytose most of the synaptic bouton (2/11 events), presynaptic material remained at the initial site confirming that microglia engage in partial phagocytosis, or trogocytosis, of synapses.

**Figure 4.**
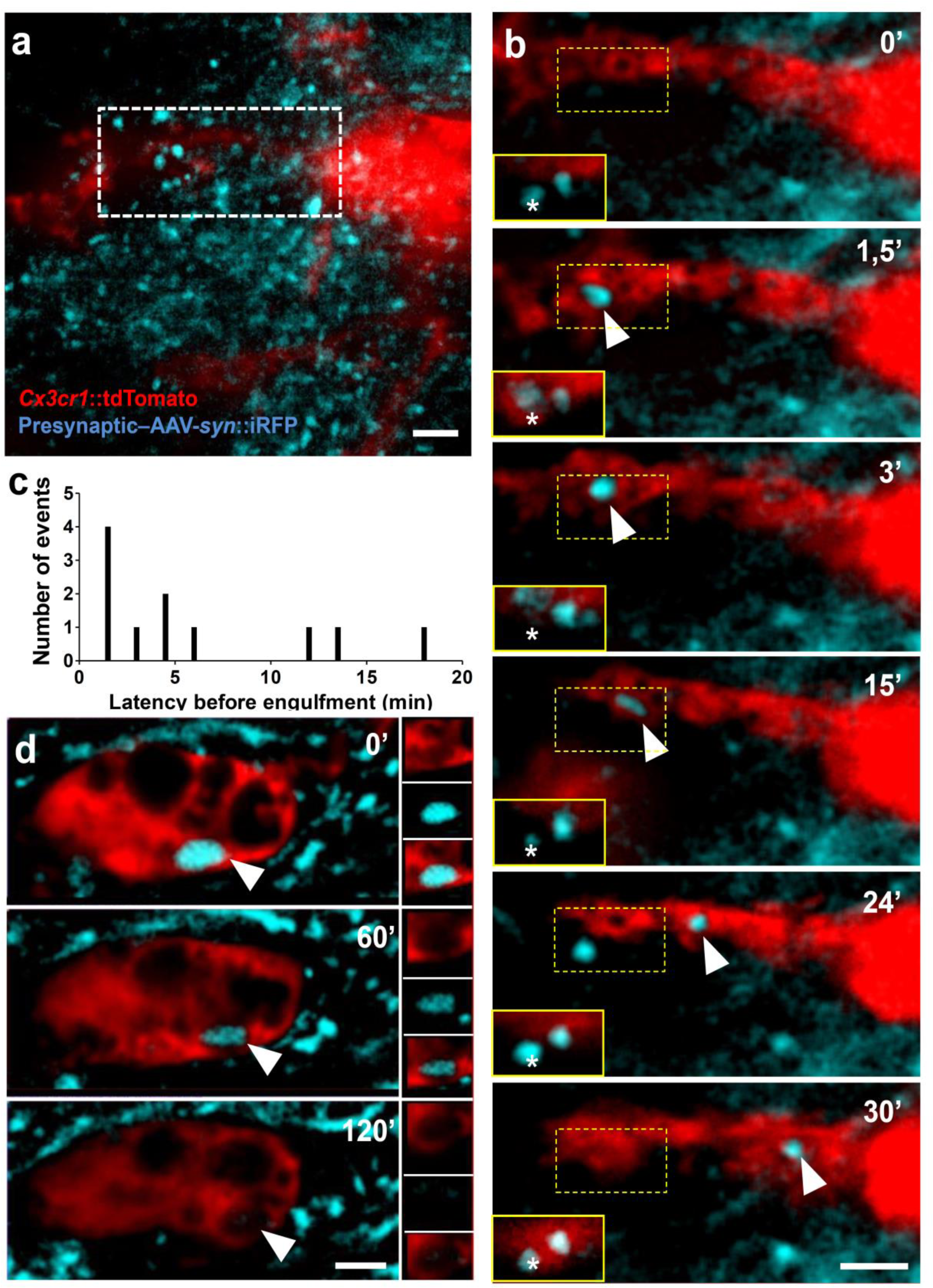
Rapid trogocytosis of presynaptic material by microglia. **(a)** Low-magnification image of a stack-projection showing microglia (red, *Cx3cr1::*CreER; *RC*::LSL-tdTomato) surrounded by ÌRFP+ presynaptic boutons from Schaffer collaterals (blue, AAV-Sy/r:iRFP) in the CA1 region of organotypic hippocampal cultures, **(b)** Time-lapse imaging revealed engulfment of a presynaptic bouton (single optical planes series of dotted box in [a]). The corresponding optical plane containing the presynaptic bouton (star) is shown in the yellow insert. While most of the bouton has been internalized by the microglia and trafficked toward the soma (arrowhead), presynaptic material remains at the original site (star), indicating partial phagocytosis, **(c)** Distribution of the latency to engulfment. **(d)** Representative image of an ÌRFP+ inclusion in a microglia soma (arrowhead) showing slow degradation (scale bar = 2 μm).

### Trogocytosis does not require CR3 signaling

The complement system has been shown to be required for the engulfment of apoptotic cells by microglia in developing hippocampus ^33^, for the efficient pruning of synapses during retinothalamic development ^6,9,34^, and for the loss of synaptic structures during neurodegeneration and aging ^35–38^. We therefore tested whether microglia trogocytosis of presynaptic elements was compromised in mice lacking the complement receptor CR3, an essential component of the complement signaling pathway expressed on microglia. Using the time-lapse imaging setup previously described, we analyzed microglia-synapse interaction in slices from C3r-KO; *Thy1*::EGFP; *Cx3cr1*::CreER; *RC*::LSL-tdTomato quadruple transgenic mice (Fig. 5a). Contrary to our hypothesis, we found no evidence for a deficit in microglia trogocytosis in knockout when compared to wild-type slices (2.3 ± 0.7 vs. 1.5 ± 0.6 trogocytosis events/cell for 3 hours respectively; p = 0.37, t-test; 6 and 8 cells analyzed from 3 cultures, Fig. 5b,c, Supplementary movie 6). There was also no difference in the latency of elimination (WT: 6.0 ± 1.6, KO: 3.9 ± 1.1, p = 0.28, t-test; Fig. 5d) or in the number of iRFP inclusions found in microglia soma (WT: 2.6 ± 0.4, KO: 2.2 ± 0.6 min, p = 0.51, t-test; Fig. 5e). These data suggest that the complement signaling pathway is not required for microglial trogocytosis of presynaptic elements.

**Figure 5.**
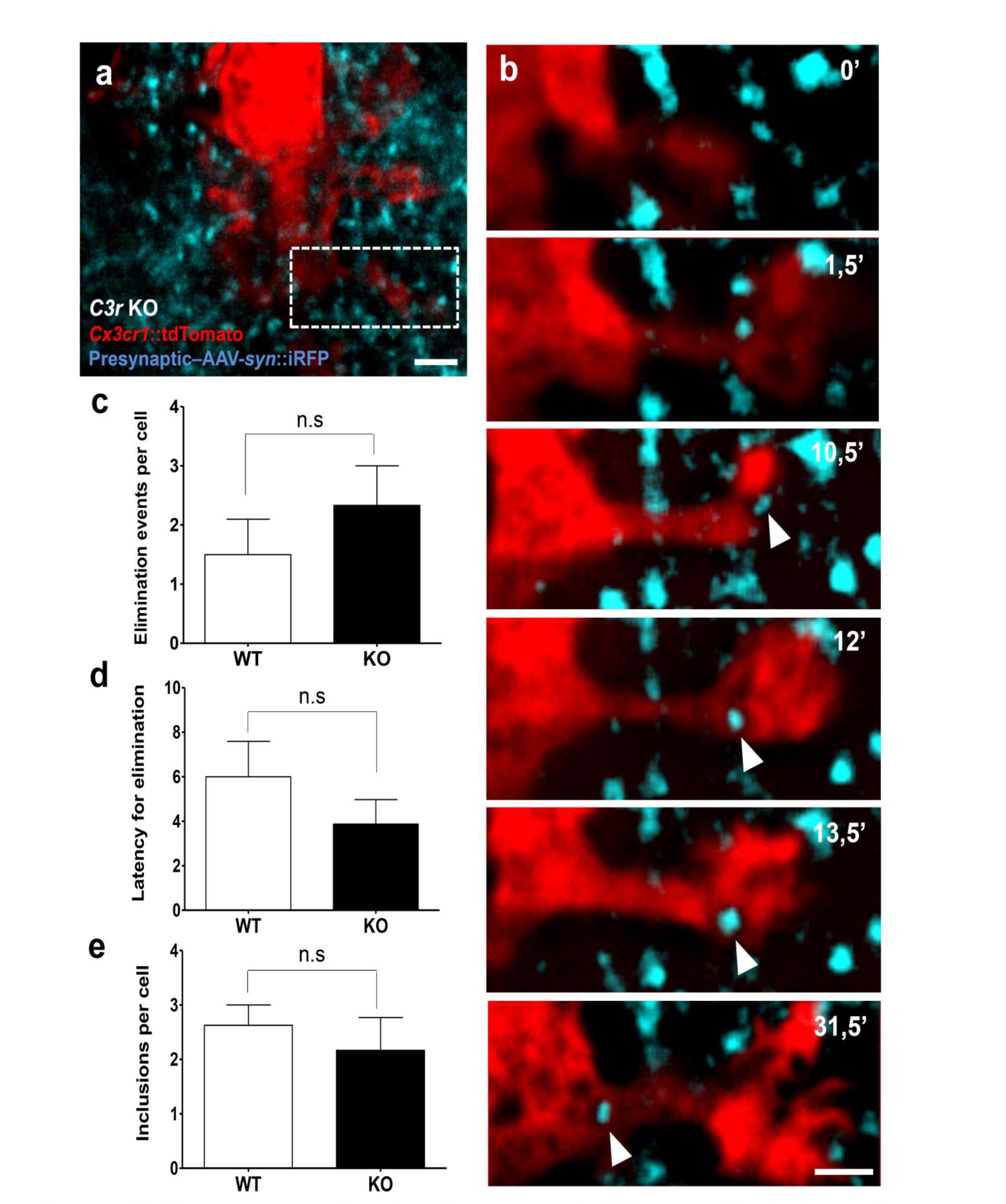
CR3 is not necessary for microglia trogocytosis. **(a)** Low-magnification image of a stack-projection showing a microglia (red, *Cx3cr1*::CreER; *RC*::LSL-tdTomato) surrounded by presynaptic boutons from Schaffer collaterals (blue, AAV-Syn::iRFP) in organotypic hippocampal slices from *C3r* KO mice, **(b)** Time-lapse imaging of *C3r* KO microglia-bouton interactions revealed engulfment of presynaptic material (from [a], dotted box). No difference was found in the **(c)** number or **(d)** latency of microglia engulfment events, or **(e)** the number of iRFP inclusions per cell between WT and *C3r* KO slices (scale bar = 2 μm).

### Microglia induce spine head filopodia formation

To explore whether microglia might indirectly induce the elimination of spines as a consequence of non-phagocytic contact or presynaptic trogocytosis we investigated microglia interactions with the postsynaptic compartment by time lapse imaging in organotypic cultures. Putative contacts between microglia processes and spines were identified and analyzed over time (Fig. 6a). Microglia-spine contacts were brief (4.2 ± 0.85 minutes) and microglia frequently re-contacted the same spine suggesting that the contacts were non-random. 10% of microglia-contacted spines (3/31) disappeared during the imaging session (Fig. 6b), and 13% both appeared and disappeared (4/31) and were classified as transient spines. However, none of these spines were in contact with microglia at the time of disappearance, arguing for a microglia-independent spine elimination process. Importantly, the frequency of disappearance of spines that had been contacted during the imaging session by microglia was not different from that of nearby (<4 μm away), non-contacted spines (10% vs. 11 %, respectively, n = 28; Fig. 6c). Transient spines, on the other hand, were found exclusively among contacted spines when compared to nearby, non-contacted spines (13% vs. 0 %). Closer, high-resolution inspection of these microglia-transient spine contact events revealed that these spines formed from filopodia that appeared at the microglia contact point or in proximity to the dendritic shaft (4/31; Fig. 4a,d,i, Supplementary movie 7) similar to a phenomenon recently observed in the mouse cortex using *in vivo* imaging^14^. Intriguingly, 39% (13/31) of the persistent, mature spines contacted by microglia formed a filopodium protruding from the head (spine head filopodia) and extending toward the microglia process (Fig. 6a,e, Supplementary movie 8), thus making spines a preferential substrate for microglia-induced filopodia compared to dendritic shaft (13 spine head filopodia vs. 4 shaft filopodia). Occasionally, we noted stretching of the entire spine during microglial contact suggesting that the microglia process was able to exert a “pulling” force on the spine head (Fig. 6a,f). Systematic, morphometric analysis of microglia-spine contact events across the imaging session was used to perform a cross-correlation analysis that revealed a significant increase in spine length that peaked just after microglia contact (n = 13 spines analyzed, Fig. 6g). Vectorial correlation of spine head filopodia and microglia process movement direction confirmed that filopodia extended toward the microglia process (n = 27 spine head filopodia formations analyzed, Fig. 6h). Notably, spine head filopodia were rarely found on nearby, non-contacted spines (7% on non-contacted spines vs. 39% of contacted spines, n = 28 and 31 respectively; Fig. 6i).

**Figure 6.**
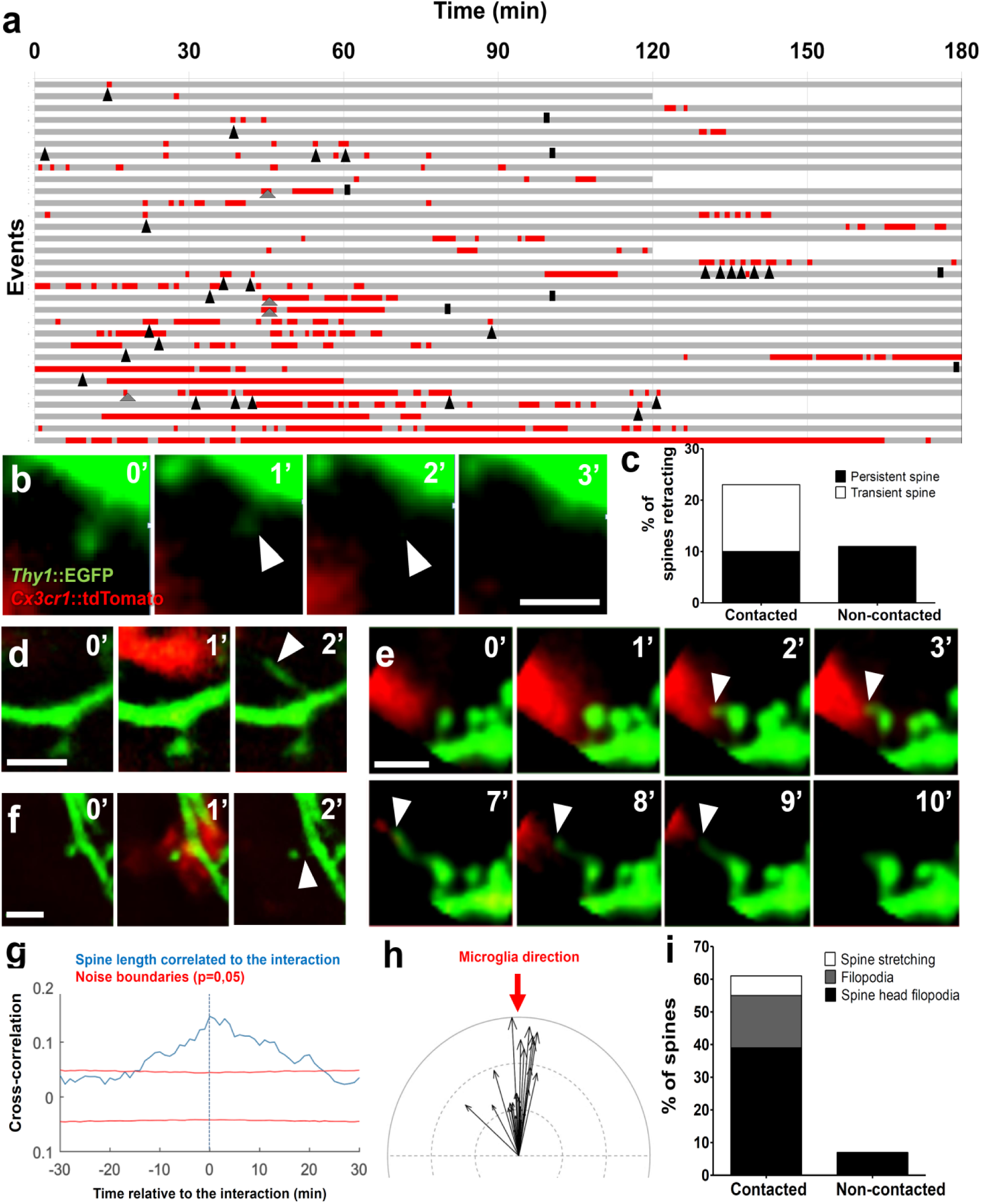
Microglia induce spine head filopodia formation. **(a)** Quantification of microglia-spine contact duration (red) over the imaging session (grey). Each line represents a spine selected to have been contacted at least once by microglia, and annotated for spine appearance (grey arrowhead), disappearance (black bar), and spine head filopodia formation (SHF, black arrowhead). Representative time sequence images of **(b)** spine disappearance, **(d)** filopodia formation, **(e)** SHF and **(f)** spine stretching, **(c)** Quantification of spine retraction rate of contacted versus non-contacted neighboring spine, **(g)** Cross-correlation analysis revealed a significant increase in spine length during microglia-spine contact, **(h)** Vectorial analysis showed a significant (Anderson-Darling test, p=0.00047) correlation of microglial process direction (red arrow) with filopodia direction (black arrows indicating filopodia length and direction, indentation = 1 μm), **(i)** Quantification of spine stretching, filopodia formation, and spine head filopodia formation events in contacted versus non-contacted neighboring spines (scale bar = 2 μm).

Spine head filopodia have been shown to contribute to the formation of new spine-bouton contacts ^39^ and proposed to be a mechanism for the movement, or “switching”, of spines from one bouton to another, possibly in response to changing synaptic activity or plasticity ^40^. Although our approach did not allow for a systematic assessment of such switching events, we did find that 27% (5/21) of microglia-induced spine head filopodia were associated with relocation of the spine head from the original site to the tip of the filopodia (Fig. 7a). Interestingly, spine head filopodia that underwent relocation showed a tendency for longer lifetimes than those that did not (27 vs. 12 min, n = 5 and 16 respectively; Fig. 7b) suggesting that this relocation might be associated with stabilizing synapse formation. This hypothesis was supported by two cases in which we were able to simultaneously image GFP+ spines and iRFP+ presynaptic boutons and we could confirm that the induced spine head filopodia made stable contact with a different, neighbouring bouton (Fig. 7c,d).

**Figure 7.**
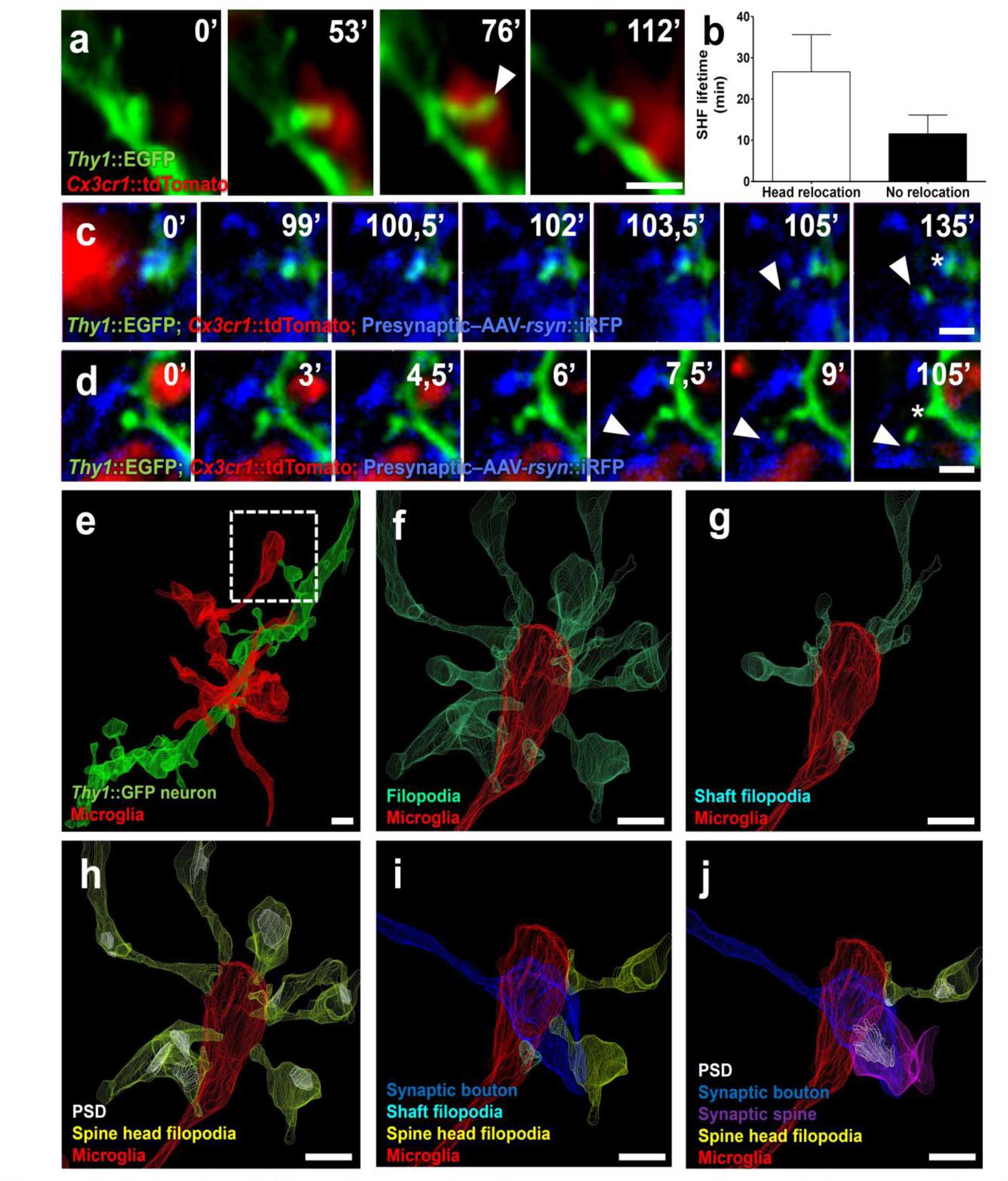
Spine head filopodia-associated synapse remodeling. **(a)** Representative time sequence images of a spine head relocation following SHF induction by microglia, **(b)** Quantification of SHF lifetime revealed more stable filopodia following relocation. **(c,d)** SHF making a stable contact with a neighboring bouton (arrowhead). In [c] the spine persists at its original location (star), while in [d] the spine relocates to the newly contacted bouton (star), **(e)** Example of microglial process in contact with a spine extending a SHF (dotted box) as identified and visualized by CLEM. Further examination of the fully segmented EM dataset revealed **(f)** multiple filopodia extending toward the microglial process, of which **(g)** a few were simple filopodia, and **(h)** the majority originated from mature spines bearing postsynaptic-densities (PSDs). Note that the microglial process is in intimate contact with a presynaptic bouton (blue) and **(i)** several of the SHFs contact the same bouton, **(j)** one of which has formed an immature PSD (arrowhead) resulting in the formation of a multiple synaptic bouton (scale bar = 1 μm).

A systematic analysis of our FIB-SEM datasets allowed us to confirm the frequent presence of microglia-associated spine head filopodia in fixed brain tissue and rule out that this phenomenon was an artifact of the *ex vivo* preparation. Remarkably, over 27% of mature, postsynaptic density (PSD)-containing spines found in contact with microglia processes presented a filopodium extending toward microglia. In one particular striking case (Fig. 7e) a microglia end-process contacting a presynaptic bouton was surrounded by converging filopodia (15 filopodia originating from 9 dendrites; Fig. 4f). Consistent with our live-imaging data, the majority (9/15) were spine head filopodia originating from mature, PSD-containing spines (Fig. 7h), while the rest were filopodia extending from dendritic shaft structures (6/15; Fig. 7g). Intriguingly, a few of these spine head filopodia extended alongside the microglia-contacted presynaptic bouton (Fig. 7i) and one appeared to initiate a synapse, resulting in the formation of a multiple synaptic bouton (MSB, Fig. 7j). We also noted that 12/15 of the converging filopodia shared their dendrite of origin suggesting that they could facilitate the formation of class I MSBs in which one bouton synapses with several spines originating from a common neuron, a form of synaptic contact that conveys increased efficacy^11^. MSB1s have previously been shown to depend on microglia-neuron signaling and contribute to the strengthening of developing hippocampal circuits and the establishment of normal brain connectivity ^11,41^. Together, these data suggest that microglia are broadly involved in structural synaptic plasticity and circuit maturation, both by the trogocytosis of presynaptic structures, as well as by the induction of filopodia from postsynaptic sites (Supplementary Fig. 3).

## Discussion

Our findings confirm the hypothesis that microglia directly engulf and eliminate synaptic material. However, contrary to previous assumptions, we found no evidence for the phagocytosis of entire synapses. Instead we observed microglia trogocytosis - or nibbling - of synaptic structures. Importantly, microglia trogocytosis was restricted to presynaptic boutons and axons, with no evidence for engulfment of postsynaptic elements. Intriguingly, microglia contacts at postsynaptic sites frequently elicited transient filopodia, most of which originated from mature spines. These data support the current hypothesis that microglia “eat” synaptic material, but point to a more nuanced role for microglia in synapse remodelling that may explain the diverse synaptic alterations observed following the disruption of microglial function.

The fact that microglia do not engulf postsynaptic structures has implications for the interpretation of previous data in the field. For example, we and others have published data showing localization of postsynaptic proteins inside microglia^8,13,16,42^. In our current confocal microscopy data we observed the intimate encapsulation of spines by microglia, a phenomenon that likely explains the previously described colocalization of immunolabelled postsynaptic proteins with microglia. However, using cytoplasmic labelling we noted that all contacted spines were still attached to the dendritic shaft via a spine neck and our threedimensional FIB-SEM reconstructions did not support the phagocytic engulfment of postsynaptic material by microglia. These data argue that immunofluorescent colocalization of postsynaptic proteins and microglia, even by super-resolution methods (e.g. stimulated-emission depletion, STED), should be interpreted with caution. Previous evidence for the localization of postsynaptic material inside microglia by single section electron microscopy, either by contrast agent-enhanced visualization of postsynaptic density (PSD)-like material^16^ or its immunodetection^8^ are potentially more difficult to counter. However, in light of our data and the limitations inherent to two-dimensional electron microscopy, caution may be warranted in the interpretation of these findings.

Our observation of microglia engulfment of synaptic material is consistent with published reports showing the localization of material deriving from axonal projections inside microglia^9^. To the best of our knowledge, our data are the first time-lapse images to directly demonstrate the active engulfment of synaptic material by microglia. Our extensive characterization of microglial content from three-dimensional FIB-SEM reconstructions validate previous data from single section electron microscopy that showed putative double-membrane inclusions of presynaptic material in microglia^9^. Moreover, our observation of phagocytic intermediates such as invaginations, pinching of presynaptic boutons and axons, and complete inclusions sheds light on the cellular mechanism involved. Synapses were previously proposed to be eliminated by microglia through phagocytosis, a process traditionally defined as the cellular uptake of particles over 0.5 μm in size^43^. Instead, our data show that only small fragments (250 nm average diameter, Fig. 3) of the presynaptic compartment are engulfed by microglia. This partial phagocytosis, or trogocytosis (from the Greek *trogo:* to nibble) has been previously described in immune and amoeboid cells^23–25^ that ingest small parts (<1 μm) of their targets within a few minutes, a timeframe compatible with our observations (Fig. 4). Although phagocytosis and trogocytosis likely share common endocytic machinery, they potentially differ in their uptake pathways. In fact we found that CR3, a microglia-expressed complement receptor involved in phagocytosis and previously proposed to mediate synapse elimination in the retinogeniculate pathway^9^, is not necessary for the trogocytosis of presynaptic structures. However, other components of the complement pathway could be involved and it is likely that different brain regions recruit different pruning pathways. One possible candidate “eat me” signal is phosphatidylserine (PS), a phospholipid known to mediate phagocytosis following its exposure on the outer leaflet of the cell membrane^44^ and recently suggested to mediate trogocytosis^25^. The known capacity of PS to laterally diffuse within the membrane could explain our observation that microglia indiscriminately trogocytose axonal shafts and boutons. Our observation that microglia trogocytosis was primarily seen at axonal shafts rather than boutons suggests that this process may be relatively non-specific and may be aimed primarily at an overall reduction in axonal processes. Alternatively, microglia trogocytosis may mediate a remodelling of axons by eliminating, for example, specific surface-associated factors that might inhibit presynaptic site formation.

Second, a major observation from our time-lapse imaging data was the induction of transient filopodia following microglia contacts. This observation was in line with a recent *in vivo* imaging study reporting that microglia contacts can induce local calcium transients in dendritic shafts followed by filopodia formation^14^. It is also consistent with a report showing that microglia participate in the learning-dependent formation of functional synapses via BDNF-TrkB signalling^15^. Together these observations argue that filopodia induction by microglia may be a widespread phenomenon and a possible trigger mechanism for the formation of functional synapses. Intriguingly, we observed that the majority of filopodia induced by microglia originate from mature spine heads (Fig. 6), a finding confirmed in our electron microscopy datasets. Our time-lapse imaging study revealed that several microglia-induced spine head filopodia formation events were followed by a relocation of the spine head to the tip of the filopodium, occasionally stabilizing at a neighbouring bouton (Fig. 7). These observations demonstrated that microglia-induced spine head filopodia can trigger spine switching. Earlier work has shown that spine head filopodia are associated with spine switching and proposed that this could allow for a rapid reorganization of synaptic networks^40^. In particular, spine switching can lead to the formation of MSBs that are formed during circuit maturation and thought to mediate the strengthening of excitatory connectivity^41^. In one particularly striking case captured in our electron microscopy study we observed over a dozen spine head filopodia converging toward a single microglia process in intimate contact with a presynaptic bouton with one of the filopodia forming a nascent MSB (Fig. 7). Although caution must be exercised when extrapolating from anecdotal electron microscopy data, the architecture of this particular case supports the hypothesis that microglia facilitate the switching of spines toward neighboring boutons and thereby promote the formation of MSBs. Such a hypothesis is consistent with previous observations that mice lacking the microglia-neuron signalling factor fractalkine (Cx3cr1-Cx3cl1) are associated with a deficiency in class I MSBs and impaired maturation of functional circuit connectivity^11^. Overall, our data argue that the prevailing view of microglia as phagocytic cells eliminating synapses during neural circuit development may be overly simplified. Instead, they suggest a broad role for microglia in synaptic remodelling via the trogocytosis of presynaptic structures and the induction and reorganization of postsynaptic sites so as to achieve an appropriate maturation of circuits.

## Materials & Methods

### Animals

C57BL/6J mice were obtained from local EMBL colonies. *Thy1*::EGFP; *Cx3cr1*::CreER; *RC*::LSL-tdTomato triple transgenic mice were obtained by crossing *Thy1*::EGFP-M^17^ (Jackson Laboratory stock 007788) with *Cx3cr1*:: creER-YFP^15^ (Jackson Laboratory stock 021160) and *Ro5a26-CSG*::loxP-STOP-loxP-tdTomato-WPRE^18^ (Jackson Laboratory stock 007905). Mice were used in homozygous state for *Thy1*::EGFP and in heterozygous state for *Cx3cr1*::CreER and *RC*::LSL-tdTomato. Cre-mediated recombination was induced by a single injection of 98% Z-isomers hydroxy-tamoxifen diluted in corn oil at 10 mg/mL (1 mg injected per 20 g of mouse weight, Sigma) at P10. Residual YFP expression in microglia yielded a faint signal in GFP channel that was thresholded out in all analysis. For Cr3-KO experiments, triple transgenic mice were additionaly crossed with CD11b-deficient mice^45^ (Jackson Laboratory stock 003991) and used in homozygous state. *Cx3cr1::*GFP mice^46^ (Jackson Laboratory stock 005582) were used in heterozygous state. All mice were on a C57BL/6J congenic background. Mice were bred, genotyped and tested at EMBL following protocols approved by the Italian Ministry of Health.

### Microglial phagocytic capacity analysis

C57BL/6J wild-type mice were anesthetized intraperitoneally with 2.5% Avertin (Sigma-Aldrich, St Louis) and perfused transcardially with 4% paraformaldehyde (PFA) at P8, P15, P28 and P40. Brains were removed and post-fixed in 4% PFA overnight (ON) at 4 °C. Coronal 50 μm sections were cut on a vibratom (Leica Microsystems, Wetzlar, Germany) and blocked in 20% normal goat serum and 0.4% Triton X-100 in PBS for 2 hours at room temperature. CD68 and Iba1 were immunodetected by overnight incubation at 4°C with primary antibodies (rat anti-CD68 1:500, Serotec; rabbit anti-Iba1 1:200, Wako) followed by secondary antibodies (goat anti-rabbit A647 and goat anti-rat A546, 1:400, Life technologies) incubation in PBS with 0.3% Triton-X100 and 5% goat serum for 2 hours at room temperature. Sections were imaged on a TCS SP5 resonant scanner confocal microscope (TCS Leica Microsystems, Mannheim) with a 63x/1.4 oil immersion objective at 48 nm lateral pixel size with an axial step of 130 nm. Microglial CD68 signal intensity was measured on Imaris software in individual cell upon 3D reconstruction using local contrast.

### Characterization of microglia-spine interactions

Brain tissue was collected at P15 as previously described. Sections were permeabilized with PBS and 0.5% triton X-100 for 30 min, and blocked with PBS, 0.3% triton and 5% goat serum for 30 min at room temperature. CD68 was immunodetected by overnight incubation at 4°C with primary antibodies (rat anti-CD68 1:500 Serotec) followed by secondary antibodies (goat anti-rat A647, 1:600, Life technologies) incubation in PBS with 0.3% triton and 10% goat serum at 4°C overnight. Secondary dendrites of bright GFP+ neurons were imaged in medial *stratum radiatum* of CA1 using Leica SP5 confocal resonant scanner microscope with a 63x/1.4 oil immersion objective, at a lateral pixel size of 40 nm and an axial step of 130 nm. Images were deconvolved using Huygens software (40 iterations, 0.1 of quality change, theoretical point spread function) and sharpened using Image J software (NIH). Interactions were determined after 3D visualization in Imaris as followed: appositions were considered when 20-50% of the spine head surface was covered by microglia, encapsulation when more than 70% was covered.

### Correlative Light and Electron microscopy

#### Confocal imaging & laser etching

Mice were perfused transcardially with PBS and fixed with 2% (w/v) PFA, 2.5% (w/v) Glutaraldehyde (TAAB) in 0.1 M Phosphate Buffer (PB) at P15. After perfusion brains were dissected and postfixed in 4% PFA in PB 0.1M overnight at 4°C. Subsequently, 60 μm thick vibratome (Leica Microsystems) coronal sections were cut and DAPI stained. Hippocampal areas were trimmed and mounted with 1% Low Melting Agarose (Sigma) in PB 0.1M on glass bottom dishes with alpha numeric grid (Ibidi). Regions of interest (ROI) containing microglia-spines interactions were imaged at high magnification with TCS SP5 resonant scanner confocal microscope with a 63x/1.2 water immersion objective, at a pixel size of 48 nm and a step size of 300 nm. Low magnification stacks containing the ROI were acquired in bright field, GFP, RFP and DAPI channels to visualize neurons and microglia together with fiducial capillaries and cell nuclei. A UV-diode laser operating at 405nm, an Argon laser at 488 nm and a DPSS solid-state laser at 561 nm were used as excitation sources. Subsequent to confocal imaging, the grid-glass bottom was separated from the plastic dish and placed onto Laser capture system microdissector microscope (Leica LMD7000) for laser etching of the ROI.

#### Sample preparation for Electron Microscopy

Upon laser branding sections were retrieved and stored in 4% PFA in PB 0.1M at 4°C. Selected sections were processed as described in *Maco et al*.^20^. Briefly, sections were washed in cold Sodium Cacodylate buffer 0.1M pH 7.4, postfixed with 1% OsO_4_/1.5% Potassium Ferrocyanide for 1h on ice, followed by a second step of 1h in 1% OsO4 in Sodium Cacodylate buffer 0.1M pH 7.4 on ice. Samples were then rinsed carefully in water and stained “en block” with 1% aqueous solution of Uranyl Acetate ON at 4°C, dehydrated with raising concentration of Ethanol and infiltrated in Propylene Oxide / Durcupan mixture with increasing concentration of resin. Durcupan embedding was carried out in a flat orientation within a sandwich of ACLAR^®^ 33C Films (Electron Microscopy Science) for 72h at 60°C. Sections were washed in cold Sodium Cacodylate buffer 0.1M pH 7.4, postfixed with 2% OsO_4_/1.5% Potassium Ferrocyanide for 1 h on ice, followed by a step with Thiocarbohydrazide for 20 min at RT and then a second step of 30 min in 2% aqueous OsO4 on ice. Samples were then rinsed carefully in water and stained “en block” first with 1% aqueous solution of Uranyl Acetate ON at 4°C and then with Lead Aspartate at 60°C for 30 min. Subsequently, sections were dehydrated with increasing concentration Acetone and infiltrated in Durcupan resin ON followed by 2 h embedding step with fresh resin. As a pilote experiment, one of the samples (Zeiss, Oberkochen, Fig. 2) was processed using a slightly modified protocol based on the application of heavy metal fixatives, stains and mordanting agents^47^ producing slightly more contrast compared to the other samples.

#### Focused Ion Beam Scanning Electron Microscopy (FIB-SEM)

Following EM processing, the flat embedded samples were trimmed to about 1 mm-width to fit on the pin for microscopic X-ray computed tomography (MicroCT). Samples were attached to the pin with either double sided tape or dental wax and mounted into the Brukker Skyscan 1272 for microCT imaging. Data was acquired over 180° at a pixel resolution of 1.5-2 μm. Karreman et al. thoroughly details the process of how the microCT data enables the correlation of fluorescent imaging to 3D EM of voluminous samples^21^. In this experiment, MicroCT revealed laser etched markings performed at the microdissector microscope, and vasculature. This vasculature, which could also be seen by negative contrast in confocal datasets, acted as fiducial features to register the various microscopy modalities (MicroCT and low and high magnification confocal data). Using Amira software (FEI Company), 3D models were generated from these microscopy modalities by thresholding and manual segmentation. These volumes could then be registered together by a manual fit to reveal the position of the event visualized by fluorescent confocal microscopy despite the loss of fluorescence during processing for EM. The registered volumes also allowed precise trimming of the sample for FIB-SEM, where it is necessary for the region of interest (ROI) to be at the surface of the sample or within 5 μm (for trimming procedure see Karreman et al.^21^). Each sample was trimmed according to the available features that would assist with later steps of FIB-SEM acquisition, such as to position the platinum deposition on the trimmed sample surface over the ROI and to position the imaging area on the cross-section face. For example, the laser markings that were made on one surface of the brain slice gave us only the ROI position in x but not over the thickness of 60 μm brain slice. For this axis, patterns made by the distribution of the vasculature were necessary to pin-point the position of the protective platinum coat over the ROI. After trimming the sample, it was mounted onto the edge of an SEM stub (Agar Scientific) with silver conductive epoxy (CircuitWorks) with the trimmed surface facing up so that it will be perpendicular to the focused ion beam (FIB). The sample was then sputter coated with gold (180s at 30mA) in a Quorum Q150RS coater before being placed in the Zeiss Crossbeam 540 focused ion beam scanning electron microscope (FIB-SEM). Once the ROI was located in the sample, Atlas3D software (Fibics Inc. and Zeiss) was used to perform sample preparation and 3D acquisitions. First a platinum protective coat of 20x20 μm was deposited with 1.5nA FIB current. The rough trench was then milled to expose the imaging cross-section with 15nA FIB current, followed with a polish at 7nA. Now that the imaging cross-section was exposed, the features visible here including vessels and nuclei were used to correlate with the registered 3D volumes in Amira and confirm the current position relative to the ROI. During the acquisition, lower resolution keyframes with a large field of view (FOV) from 40x40 to 70x70 μm were acquired in order to have this broader context of the sample. Provided there were enough features close to the ROI, this information helped to position the high resolution imaging field of view (typically 10x10 μm). The 3D acquisition milling was done with 3nA FIB current. For SEM imaging, the beam was operated at 1.5kV/700pA in analytic mode using the EsB detector (1.1kV collector voltage) at a dwell time from 6-8 μs with no line averaging over a pixel size of 5x5nm and slice thickness 8nm. For the pilot acquisition run at Zeiss, Oberkochen, a large volume was first acquired at low magnification without prior microCT while correlating with the confocal dataset to detect the ROI, which was subsequently imaged at 5 nm isotropic pixel size.

#### Image processing and segmentation

A single stack file containing individual FIB images was aligned on ImageJ software (https://imagej.nih.gov/ij/) with the help of the linear stack registration plugin (SIFT). Grayscale look up table was inverted and the stack was binned 2X in both lateral and axial planes. Microglia and dendrites of interest were located based on their *xyz* coordinates within the ROI and from fiducials correlated between EM and confocal datasets. The segmentation was carried out manually using iMOD software (http://bio3d.colorado.edu/imod/) and 3D model was generated and matched to the confocal dataset to confirm perfect correlation. Blender (https://www.blender.org/) was used as software to generate 3D animation of the full EM dataset combined with the 3D model.

### Live imaging of hippocampal slice culture

#### Preparation of the slices

Organotypic hippocampal slice cultures were prepared using the air/medium interface method^48^. Briefly, mice were decapitated at P4 and hippocampi were dissected out in cold dissecting medium (HBSS 1x (Gibco), Penicillin/streptomycin 1x, HEPES (Gibco) 15 mM, Glucose (Sigma) 0.5%). Transverse sections of 300 μm thickness were cut using a tissue chopper. Slices were laid on culture-inserts (Millipore) in pre-warmed 6-wells plates containing 1.2 mL of maintaining medium (MEM 0.5x (Gibco), BME 25%, Horse serum 25%, Penicillin/streptomycin 1x, GlutaMAX 2 mM, Glucose 0.65%, Sodium Bicarbonate 7.5%, ddH2O qsp). Medium was replaced after 24 hours and then every 2 days. For culture of *Thy1*::EGFP; *Cx3cr1*::CreER; *RC*::LSL-tdTomato brain slices, 98% Z isomers-OHT was added to the maintaining medium at 0.1 μM during the first 24 hours. After preparation, cultures were maintained for up to 21 days in vitro (DIV21) in incubator at 35°C and 5% CO2. The morphology of microglia was inspected in Cx3cr1::GFP slices at the indicated time-points upon fixation in PFA 4% for 1 h at room temperature, PBS-washed, mounted with mowiol, and imaged on a Leica SP5 confocal resonant scanner microscope with a 63x/1.4 oil immersion objective.

#### Light sheet live-imaging

Live-imaging of microglia and synapses in hippocampal slice cultures (DIV10-19) was performed using a Z1 light sheet microscope (Zeiss). The imaging chamber was set at 35°C and 5% CO2, and filled with imaging medium (MEM without phenol red 0.5x, Horse serum 25%, Penicillin/streptomycin 1x, GlutaMAX 2 mM, Glucose 0.65%, Sodium Bicarbonate 7.5%, ddH2O qsp) 2 hours prior the imaging session to allow the system to equilibrate and the medium to reach pH7. Low melting point agarose was prepared at 2% with imaging medium and incubated at 35°C and 5% CO2 for 30 min for reaching proper pH. The membrane of the Millipore insert containing the slice of interest was cut around the slice, and laid onto the equilibrated liquid agarose before polymerization at 4°C for 1 min. The slice mounted on agarose was then placed in incubation at 35°C and 5% CO2 for further 30 min. The polymerized agarose containing the slice was then inserted into a FEP tube adapted on a glass capillary, and the slice was gently pushed to be exposed 1 mm outside the FEP tube. The capillary was then placed on the microscope holder, and the slice was immersed in the imaging medium of the chamber 1 hour prior imaging for stabilization. For all imaging sessions, microglia and neurons were selected for their brightness and position in the *stratum radiatum* of CA1, 2 to 30 μm from the slice surface. Imaging was performed for 2-3 hours using a 60x/NA1 water immersion objective, with a lateral pixel size of 130 nm and an axial step of 480 nm. For microglia-postsynaptic structures interactions (2-colors imaging), 488 and 561 lasers were used for simultaneous acquisition of GFP and tdTomato signal using 505-545 and 575-615 band pass filters on 2 cameras, at a rate of 1 frame/45-60 seconds. For microglia-pre/postsynaptic structures interactions (3-colors imaging), 2 channels were fast switching between frames: 488 and 561 lasers were used for simultaneous acquisition of GFP and tdTomato signal using a 505-545 band pass filter and a 585 long pass filter, and iRFP was imaged using a 638 laser line and the 585 long pass filter, at a rate of 1 frame/90 seconds. All datasets were deconvolved using Zen software, and corrected for drifts on Image J using a script created by Albert Cardona and Robert Bryson-Richardson^49^ and modified by Cristian Tischer (EMBL Heidelberg).

#### Analysis of microglia motility

TdTomato signal intensity was measured in microglial processes and normalized across all datasets. Noise was measured outside microglia, and removed by thresholding to the measured value + 40%. Motility was assessed by analyzing protrusions over one minute interval. Extending and retracting protrusions were counted and measured at 0, 60, and 120 min after the beginning of the session to confirm imaging.

#### Fluorescence labeling of presynaptic structures

AAV-*rSyn*::iRFP670 virus was generated by cloning AAV vector serotype 2 ITRs with a rat Synapsin promoter (a gift from Hirokazu Hirai^50^), a iRFP670 coding sequence (a gift from Vladislav Verkhusha^51^, Addgene plasmid # 45457), WPRE and human growth hormon polyA sequence. Viral production and purification was performed according to McClure et al^52^. with minor modifications. Briefly, 15 x 15 cm dishes of HEK293T cells were transfected with the pAAV-rSyn::iRFP670 plasmid, together with pAAV1, pAAV2 and the helper plasmid pFdelta6 using PEI (Sigma, 408727). 72 hours after transfection the cells were collected and lysed according to the protocol and the virus containing-cell lysate was loaded onto HiTrap Heparin columns (GE Biosciences 17-0406-01). After a series of washes, the virus was eluted from the heparin column to a final concentration of 450 mM NaCl. Finally, the virus was concentrated using Amicon Ultra-15 centrifugal filter units (Millipore UFC910024) and semiquantitative titering of viral particles was executed by Coomassie staining versus standards. CA3 neurons of hippocampal slices were infected with AAV-rSyn::iRFP670 the day following the preparation of the culture, by a local injection of 0.1 μL of virus in the pyramidal layer of CA3 using a glass capillary under a stereomicroscope. Expression of iRFP was observed in CA3 neurons exclusively as early as 5 days post infection, and expression in CA3-CA1 Schaffer collaterals reached satisfactory expression level 10 days after infection.

#### Analysis of microglia-presynaptic compartment interactions

Contacts between microglia and boutons were detected by scouring iRFP+ axons in 3-dimensions for the entire duration of the imaging session. Elimination events were defined as clear engulfment of iRFP+ material by microglia from an iRFP+ bouton. Latency before engulfment was measured as the time of contact between microglia and the bouton before any iRFP+ material was seen internalized. Inclusions were defined as iRFP+ structures within microglia for which the origin could not be defined as they were present from the beginning of the imaging session.

#### Analysis of microglia-postsynaptic compartment interactions

Contacts between microglia and the postsynaptic compartment were detected by scouring dendritic shafts of GFP+ neurons in 3-dimensions for the entire duration of the imaging session. All z-planes containing the contacted spine over time were axially projected. GFP intensity was measured in the dendritic shaft and normalized across all the datasets. GFP signal noise and residual microglial YFP signal were measured in microglial processes, and throsholded out to the measured value + 40%. For all spines that were seen to be contacted at least once by microglia, we measured the spine length, the size of the head, and the extent of contact between the spine and microglia for each timepoint of the entire imaging session. The length of the spine was measured from the dendritic shaft to the tip of the spine head, the change in head size was measured as variation in GFP signal at the spine head, and the extent of contact was measured as the percentage of GFP signal at the spine head colocalizing with tdTomato signal. We then performed cross-correlation analysis over time of the extent of contact, with the spine length variation (correlation with the head size not shown) using bootstrap resampling on MATLAB.

#### Statistical analysis

All data are represented as mean ± SEM. To determine statistical significance, data distribution was first tested for normality, and the corresponding t-test (parametric or non-parametric) was performed using the Graphpad Prism software. For multiple group analysis (motility analysis over time) two-way ANOVA was performed using Graphpad Prism software. Cross-correlation analysis was made using MATLAB software.

## Acknowledgements

We would like to thank Hiroki Asari for his help with the cross-correlation analysis using MATLAB, Andreas Buness for vectorial correlation on MATLAB, Tom Boissonnet for generating the CLEM animation with Blender, Senthilkumar Deivasigamani for careful reading of the manuscript and critical input. Funding was provided by EMBL (C.T.G., L.W.), ERC Advanced Grant COREFEAR (C.T.G), and the People Programme of the European Union's Seventh Framework Programme FP7/2007-2013/ under REA grant agreement n°327409 (U.N.).

## Competing interest

The authors declare no competing interest.

## Author contributions

L.W. and C.T.G. designed the experiments. L.W. performed the experiments and analyzed the data, except as follows. U.N. and A.V. performed the analysis of microglia phagocytic capacity. P.M. and N.S. performed FIB-SEM acquisition under the supervision of Y.S. and G.B. G.dB. performed the microglia motility analysis. M.E. performed microglia-presynaptic interactions analysis. A.R. produced the AAV-*rSyn*::iRFP virus. A.S. performed pilot FIB-SEM acquisition. L.W. and C.T.G wrote the paper.

